# Laser-based killing of a macroparasite inside its live invertebrate host

**DOI:** 10.1101/2023.03.06.531243

**Authors:** Olivier Musset, Aude Balourdet, Marie-Jeanne Perrot-Minnot

## Abstract

Whether host phenotypic alterations induced by parasites are reversible or not is a core issue to understand underlying mechanisms as well as the fitness costs of infection and recovery to the host. Clearing infection is an essential step to address this issue, which turns out to be challenging with endoparasites of large size relative to that of their host. Here, we took advantage of the lethality, contactless and versatility of high-energy laser beam to design such tool, using thorny-headed worms and their amphipod intermediate host as a model system. We show that laser-based de-parasitization can be achieved using blue laser targeting carotenoid pigments in *Polymorphus minutus* but not in the larger and less pigmented *Pomphorhynchus tereticollis*. Using DNA degradation to establish parasite death, we found that 80% *P. minutus* died from within-host exposure to 5 pulses of 50 ms duration, 1.4W power and 450 nm wavelength. Survival rate of infected gammarids to laser treatment was higher than uninfected ones (62% and 33% at 11 days after exposure, respectively). The failure to kill *P. tereticollis* was also observed with nanosecond-green laser, an alternative laser source targeting lipid. We discuss the possible causes of amphipod death following parasite treatment and highlight the perspectives that this technology offers.

## Introduction

Host-parasite interactions are a driven force in the ecology and evolution of species and, as such, are extensively studied. Standard methodology to address the molecular, physiological, behavioral, and life-history responses of hosts to infection relies on the comparison of naturally or experimentally infected hosts to uninfected ones. In addition, curing host from infection allows addressing the reversibility of host responses, as well as assessing fitness costs associated with the parasite’s strategy of host exploitation. This is generally achieved by pharmaceutical treatment in humans and in domesticated or captive animals. Alternatively, the technology of lasers could offer promising alternatives to clear parasites because it offers a contactless tool to deliver lethal high-energy beam and could be highly specific. It has been implemented for a wide range of usage in medical sciences (with a dedicated journal, Lasers in Medical Science), but its application to non-human species is much less developed. Recent applications are limited and include short laser shot of various wavelengths to target vector-born parasites by killing mosquitoes (Keller et al. 2016) and an automated 532 nm laser-beam to kill salmon lice (ectoparasitic copepods) in rearing tanks (Stingray® patent EP2531022, Stingray Marine Solutions AS, Norway). In both cases however, the targets (parasite vector and ectoparasite) were external and readily exposed to a laser beam. Targeting endoparasites without killing the host is more challenging; it has been restricted so far to killing microparasites, using conjugates of gold nanoparticles and antibodies and subsequent laser irradiation (Pissuwan et al. 2007). Yet it could also be used to selectively damage macro-endoparasites within host body, as far as the parasite can be selectively targeted and concentrate most of the energy delivered to limit deleterious effects on the surrounding host tissues.

Laser-directed optical energy deposition is almost instantly converted into heat. The induced heating is then distributed temporally and spatially in the material according to its thermal properties, for example conductivity or thermal diffusivity. Depending on the type of laser used, irradiation can induce thermal effects such as heating, fusion or vaporization, photochemical effect (molecular denaturation) or photo-mechanical effect (fast thermal ablation) that leads to the expulsion of material from the exposed substrate. The thermal effects correspond to the formation of a thermal gradient of varying duration and possibly to one or more phase changes in the material. They are based on optical absorption as described by Beer-Lambert’s law, and on local transformation of absorbed light into heat. The induced heating is then distributed temporally and spatially in the material according to its thermal properties, for example conductivity or thermal diffusivity. In general, thermal effects can be divided into two main categories according to the laser exposure time. Rapid effects correspond to an almost instantaneous supply and transformation of energy into heat in the material and a long-lasting heat propagation. Slow effects occurs when heat generation time and propagation time are of the same order of magnitude. In addition to absorption, the propagation of light in soft tissues is also guided by diffusion in a tissue-specific manner. The diffusion phenomenon reduces the density of laser energy deposited in target tissue upon absorption and causes the angular spreading of the incident laser light to non-target surrounding tissues. The diffusion phenomenon and consecutive collateral damages can be minimized by using a wavelength matching the optical absorption properties of the target tissues.

Our aim was to develop a method to kill a macro-endoparasite in vivo while preserving host viability, taking advantage of the great versatility of laser-based technology. Our host-parasite system involves the freshwater crustacea *Gammarus fossarum* as the intermediate host of two species of thorny-headed worms (Acanthocephala): *Pomphorhynchus tereticollis* and *Polymorphus minutus*. The last larval stage of the parasite, called cystacanth, appears as an opaque ball yellow to red in color, oval-shaped, and ranging from 0.25 to 0.65 mm^3^ in volume depending on the species. In cystacanths of *P. minutus*, lipids contribute to 28% of dry weight, most of them being neutral lipids deposited in the radial layer of the body wall (Crompton, 1970). Carotenoid pigments are stored in the outer tegument of the cystacanths of both acanthocephalan species [3,4]. The amphipod *G. fossarum* is approx. 0.5 to 0.8 cm length and hosts one to several cystacanths in its body cavity (Plate S1). It is visible through the translucent cuticle of the host, at least over a good part of the visible and near infrared light spectrum (Kaldonski et al. 2009).

We tested two laser sources corresponding to either slow or rapid effects as described above, and matching either the parasites’ chromophore responsible for their integumental pigmentation or the constitutive molecules of their tegument (carotenoids and lipids, respectively; Crompton, 1970). Their features can be summarized as follows: (1) A continuous laser source at the absorption peak of carotenoids (450nm) [Gaillard et al. 2004; Kaldonski et al. 2009), with output power around the watt, possibly chopped, and an exposure time of the order of a second to a few seconds. The cuticle was first drilled by vaporization at low power, and then thermal heating at high power was expected to induce lethal damage to the parasite. (2) A pulse laser source matching absorption peak of lipids constitutive of acanthocephalan cystacanth at 532 mn, with a very short pulse duration typically in the nanosecond range (<100ns), low pulse energy (<10mJ) and a low repetition frequency (<100Hz). We expected rapid material removal by thermal ablation, i.e., a mechanical wave induced in the tissue. The expected effects of these two laser treatments were essentially thermal but differed for the two laser sources as suggested by their kinematics, the induced thermal gradient of varying duration and possibly one or more phase changes in the material. The efficiency of these methods was assessed by observing the mechanical and thermal damage done at the organism level, and the survival of both the targeted cystacanth and its gammarid host.

## Material and methods

### Guiding principles relating to the design of the experiment and full set up, and biological material

General background information on laser - material interactions and the physical properties of lasers are necessary to guide specific methodological choices and assess their applicability to de-parasitization. Given the paucity of information on the biological material itself (acanthocephalan tissues) and therefore on the behavior of the laser beam during its propagation in its target (cystacanth) and outside (host body), we based our approach on theoretical models of light – energy transformation and of the spatial profile of laser beam according to its optical shaping, as detailed in electronic supplementary material, S1. These allowed us to predict the performance and features of the two laser sources chosen - Blue-Diode Laser, BDL, at 450 nm, and Nanoseconds-Green Laser, NGL, at 532 nm -, including laser spot diameter, M^2^ beam quality, output energy, fluence and peak illumination (electronic supplementary material S2). The principles guiding the choice of laser sources and the design of optical shaping system as well as the full setup in details, are presented in electronic supplementary material (S1, S2). The two laser sources and their operating system were incorporated to a full setup comprising the necessary elements to visualize and immobilize the individual gammarid and focus the laser-beam on its target, the parasite. The optimization of laser treatment was reached by testing different combinations of pulses number and duration per shot.

All gammarids *G. fossarum* were collected in River Norges at Orgeux (Burgundy, France: 47°21’40.61” N, 5°9’30.53’’ E). Gammarids were allowed to acclimate for at least one week prior to experiments, in tanks filled with oxygenated dechlorinated ultraviolet (UV)- treated tap water mixed with water and rocky substrate from the river (‘conditioned’ water, hereafter abbreviated as CW). Prior to and after laser treatments, gammarids were maintained in a temperature- and photoperiod-controlled room (16°C, UV, 12 L: 12 D, under 800 Lux illumination), and fed on decaying elm leaves and occasionally commercial blood worm.

### Laser sources: Blue-Diode Laser and Nanosecond-Green Laser

We used a high-power multimode laser diode capable of emitting 1.6W @ 450nm (PL-TB450B OSRAM) with an M^2^ beam quality factor of about 4. As desired, BDL has its emission wavelength within the absorption maxima of carotenoids. The laser beam at diode output was originally divergent and astigmatic, with strong divergences in the two main directions orthogonal to the direction of propagation. We added several optical components at the output of BDL to generate a laser beam of diameter as small as less than 100μm with a working distance to sample around 100mm and a millimeter Rayleigth distance (see electronic supplementary material S2 for details). The BDL was controlled by an OEM driver from Thorlabs. A low current mode of operation allows the power diode to be used directly as an aiming beam. A simple Arduino Due board with a color touchscreen provided a user-friendly means of setting the parameters of laser shot: the energy, duration and number of pulses, and the choice of up to 5 consecutive shots with different specifications.

The NGL source was used for thermomechanical effects (electronic supplementary material S1). This lab-made laser converted the natural emission wavelength of 1064nm - in the near infrared - into green emission at 532nm with a second harmonic generator, to obtain pulse energy from 1 to 20mJ and pulse duration around 10 ns with a maximum repetition frequency of 40Hz. Several laser’s parameters such as firing frequency, the number of pulses and energy per pulse were adjustable. The laser beam quality described by the parameter M^2^ was less than 2. In this field of intensity, the interactions were highly photomechanical with the possible appearance of a plasma.

### Full Setup

The full setup was designed to configure and operate the two laser sources and to pilot sample holder from a single control box. It should enable to switch from one laser source to another easily and quickly, and to be able to precisely position the laser beam on the target area (with step less than 50μm). The setup included the two pulsed and continuous laser sources with their optical shaping and focusing systems, a thermo-regulated sample holder mounted on a motorized XY displacement (precision of 50 μm; Thorlabs LTS25M-Z8 for displacements, KDC101 for the controllers), and a lighting and aiming system including a color camera (Dino-Lite Premier) (Fig. 1).

**Figure 1.**
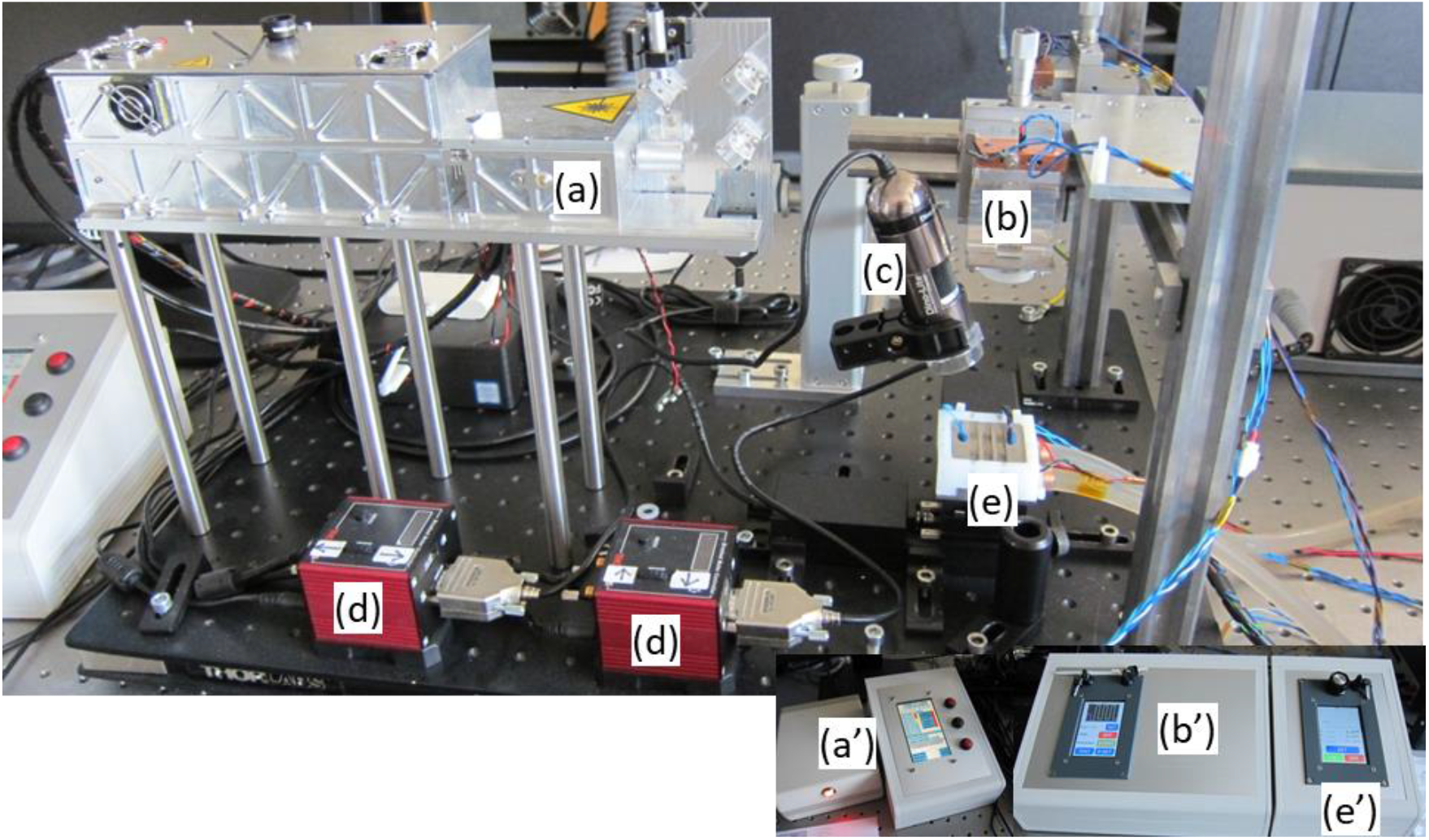
Complete setup for irradiating acanthocephalan macroparasite inside live crustacean host, comprising two laser sources – Nanoseconds-Green laser (532 nm) (a) and Blue-Diode laser (450 nm) (b) -, a video camera (c), the 2D-displacement system of temperature-controlled sample holder (d, e) (connected to computer screen, not on the picture); (a’, b’, e’): control units of BDL and NGL, and of refrigerated sample holder (thermostat), respectively.

The thermo-regulated sample holder was specifically designed (in addition to an anesthetic procedure, see below) to fully immobilize gammarids and minimize heating during irradiation. The sample holder comprised several stainless-steel channels to allow easy manipulation of gammarids and was designed for precise alignment of the laser spot on the parasite inside its host. Complete immobility of gammarids during laser treatment was mandatory to target the cystacanth in a host that is small-sized relative to the parasite, and hence to limit possible damages to host’s surrounding tissues. Immobility also prevented stress-sensitive reactions to light, handling, and lack of oxygen. We therefore combined cold and anesthesia to immobilize gammarids, by thermo-regulating sample holder at a stabilized temperature of 3°C, and bathing gammarids in the anesthetic ms222 at 60 mg.L^-1^ for approx. 30 min. prior to laser treatment (Perrot-Minnot et al. 2021). A detailed description of the overall setup (power supplies, steering system, aiming system), is provided in electronic supplementary material (S2). After irradiation, gammarids were maintained for no more than 2 hours in a portative ice box refrigerated at 10-15°C., and then transferred and maintained in groups of 10 for maintenance.

### Monitoring the death of *P. minutus* and *P. tereticollis*, and the survival of infected and uninfected gammarids to laser treatment

Two proxies were used to assess the survival of *P. minutus* and *P. tereticollis* cystacanths to laser irradiation: cystacanth’s capacity to evaginate the proboscis (Perrot-Minnot et al. 2011), and DNA integrity. The proboscis is a spiny structure invaginated in the cystacanth, used for attachment to the intestinal wall of the definitive host (bird or fish) after predation of infected gammarids by a predator final host; its evagination is metabolically costly and expected to occur in healthy cystacanths in response to a stimulus (bile from fish or elevated temperature for acanthocephalan of fish and bird, respectively). The level of DNA integrity is expected to be a reliable indicator of death, since high molecular weight DNA is rapidly degraded by endogenous exonucleases, leading to DNA fragmentation short time after cell death (Roy et al. 2020).

Evagination assay was performed just after dissection of the infected gammarid at least 24 hs after irradiation, by immersion in the dark of *P. minutus* cystacanth in ddH20 at 42°C and of *P. tereticollis* cystacanth in bile from barbels at 16°C. Under these conditions, evagination occurs within 1 to 1.5 hs (Perrot-Minnot et al. 2011). We assessed the integrity of total DNA from deep-frozen cystacanth, dissected out the host at least one day and up to one month after laser exposure. DNA extraction from single cystacanth was restricted to digestion without solvent extraction and subsequent precipitation, as these steps could decrease extraction yield and are not required for the analysis of DNA fragmentation (Park and Patek, 1998). RNAse treatment was not systematically done, as preliminary tests shown that, given the low amount of total DNA in a single cystacanth, DNA degradation was not evidenced in a visible smear but only in the disappearance of large molecular weight DNA. Details on DNA digestion, concentration by freeze-dried lyophilization, and visualization by agarose gel electrophoresis are given in electronic supplementary material (S3).

Survival of exposed gammarids was monitored during eleven days in air-conditioned room at 16°C. Gammarids were pooled by 8-10 individuals in 10.5 * 16 cm rectangular boxes filled with UV-treated and filtered tap water mixed with water and rocky substrate from the Norges River and were provided with decaying elm leaves and chironomid larvae as food, and rocks as refuge.

### Statistical analysis

All analysis were done with R-Studio, version 1.3.1073 (RStudio Team, 2020).

Survival of gammarids to laser treatment was analyzed using the Cox proportional-hazards regression model with survival time as dependent variable and laser treatment, infection status (infected or uninfected) and the interaction of both as predictors (‘survival’ package v. 2.44-1.1; Therneau 2021). We computed Type-II likelihood-ratio tests for each covariate in the model using Anova function in the ‘car’ package v. 3.0-11 (Fox and Weisberg 2019) We also checked model assumption of proportional hazards by visual inspection of scaled Schoenfeld residuals against time, for each covariate.

We compared the proportion of evaginated cystacanth in each laser treatments to the unexposed control the using Fisher exact test on contingency table (Package stats v. 4.1.2). Odds ratio of evagination and associated 95% CI were estimated for each laser treatment. We analyzed the proportion of cystacanth showing DNA integrity similarly.

## Results

The setup of the BDL and additional optical components delivered a laser beam of 1.4W with a minimum spot size was of around 70×30μm (≈0.002 mm^2^) corresponding to an irradiance of 67kW/cm^2^ (Table 1). Energy distribution at the focal point is presented in electronic supplementary material S2 (Fig. S3). The exposure of parasite inside host body was facilitated by pre-drilling a hole through the cuticle with a laser beam of 0.5W for a duration of 100ms, equivalent to an energy of 50mJ (area of pre-drilled hole < 0.008 mm, N=4). The area of laser-induced wound on gammarid’s cuticle moderately increased after a subsequent shot of 1.4W for 100 ms, equivalent to an optical energy of 140mJ (mean area < 0.02 mm^2^, N=7). The setup of the NGL and additional optical components delivered a laser beam at 532 nm with energy per pulse up to 5mJ, corresponding to a peak irradiance of up to several GW/cm^2^ for a 10ns pulse duration (Table 1).

**Table 1.** Summary of the various laser treatments tested to kill acanthocephalan cystacanth inside their live amphipod host, using two laser sources (Blue-Diode laser and Nanoseconds-Green laser).

| Type of laser | Wavelength (nm) | Beam size @focus spot with 100 mm focal lens | Pulse duration (s) | Power (W) | Peak Intensity (kW.cm<sup>-2</sup>) | Fluence per pulse (kJ.cm<sup>-2</sup>) |
| --- | --- | --- | --- | --- | --- | --- |
| Chopped CW Blue-Diode laser | 450 | 70 x 30µm | 20.10-3 | 1.4 | 67 | 1.3 |
|  | 450 | 70 x 30µm | 50.10-3 | 1.4 | 67 | 3.3 |
|  | 450 | 70 x 30µm | 100.10-3 | 0.5 | 24 | 2.4 |
|  | 450 | 70 x 30µm | 100.10-3 | 1.4 | 67 | 6.7 |
| Nanosecond s-Green Q-switched laser | 532 | ≤ 50µm | 10.10-9 | Up to 500kW | Several GW | Few tens to few hundreds |

### Efficiency of laser treatment with the Blue-Diode Laser on *P. minutus*

Cystacanths of *P. minutus* exposed to a laser beam of 5 pulses of 50 ms at 1.4W exhibited clear wounds in the form of protuberance or lesion (Fig. 2). The evagination success of *P. minutus* was dependent on laser treatment, with a success rate of unexposed cystacanths, cystacanths exposed to 3 pulses of 20 ms, and cystacanths exposed to 5 pulses of 50 ms of 91% (N=22), 27.3% (N=22) and 4% (N=25), respectively. The Odd ratios of failure to evaginate after laser irradiation compared to controls were high for both laser treatments and was more than seven times higher at 5*50 ms than at 3*20 ms (OR = 177.1 [16.8 - 9388.9], *P* < 0.0001, and OR = 24.1, 95%CI [4.03 - 273.5], *P* < 0.0001, respectively).

**Figure 2.**
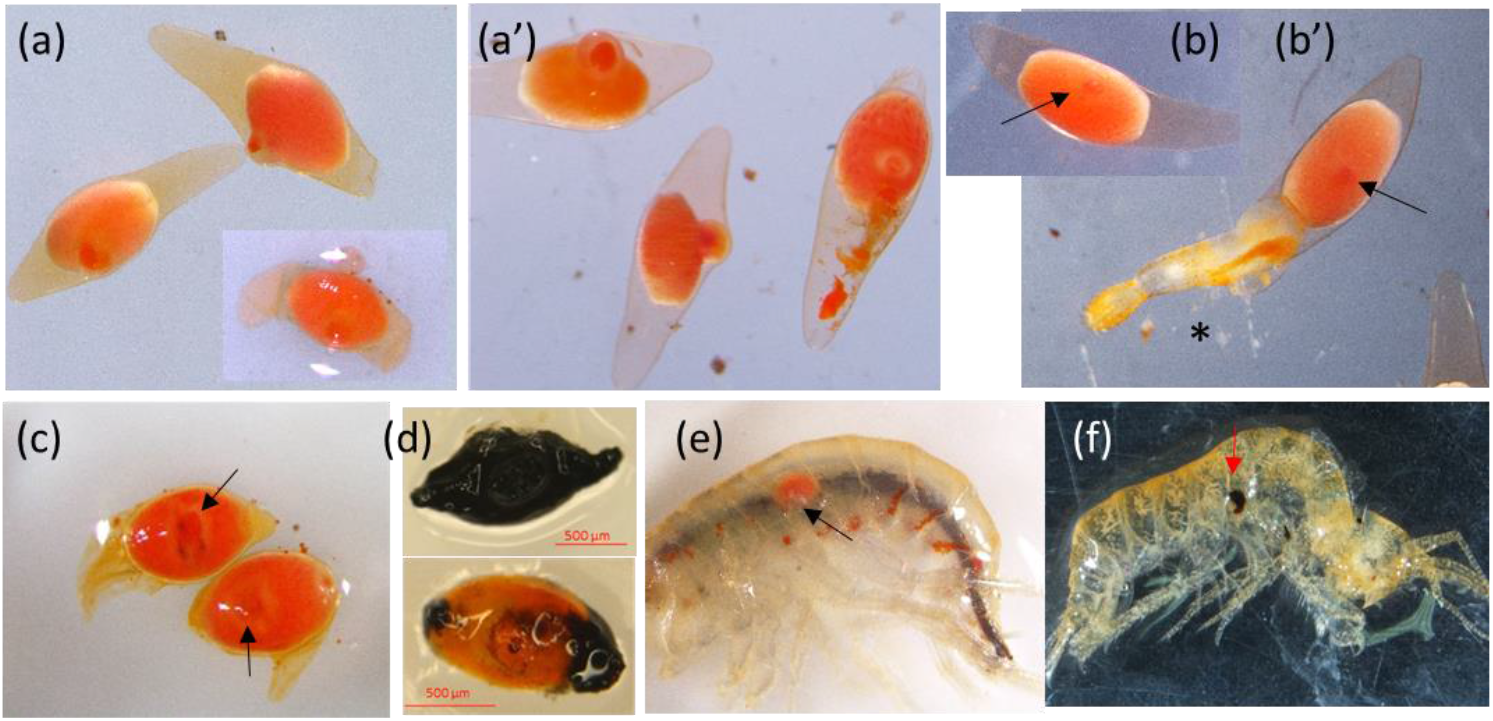
Appearance of *P. minutus* cystacanths (larval stage) dissected out gammarid host after exposure to Blue-Diode laser at 450 nm 1.4W for 3 pulses of 20 ms (a, a’, b, b’) or 5 pulses of 50 ms (c, d). After exposure at 3 pulses of 20 ms, the same cystacanths were observed before (a, b) and after (a’, b’) evagination assay (42°C, 1 hr): (a’) failure to evaginate, (b’) evagination of trunk and proboscis (*). After exposure at 5 * 50 ms, cystacanths were observed either few hours after irradiation (c) or a month later showing complete (top) or partial (bottom) melanization on exceptional occasion (d). (e, f) Live *G. fossarum* infected with *P. minutus* several days after laser treatment (e) and its exuviae after molting (f): the melanized scar is no more visible on the live gammarids (e, black arrow), as it has gone with the exuviae upon molting (f, red arrow) (gammarids length is 6 to 10 mm).

DNA integrity assay revealed the complete degradation of DNA in 50 out of 63 cystacanths exposed to 5*50 ms (80%) (Fig. 3a,b,c), most of them analyzed one month after laser exposure. The large molecular weight genomic DNA was not visible anymore in dead cystacanths, whereas small DNA fragments remained visible (Fig. 3). By contrast, DNA integrity (and RNAs in absence of RNAase treatment) was preserved in cystacanths exposed to 3*20 ms (N=9) and in unexposed controls (N=38) (Fig. 3a).

**Figure 3.**
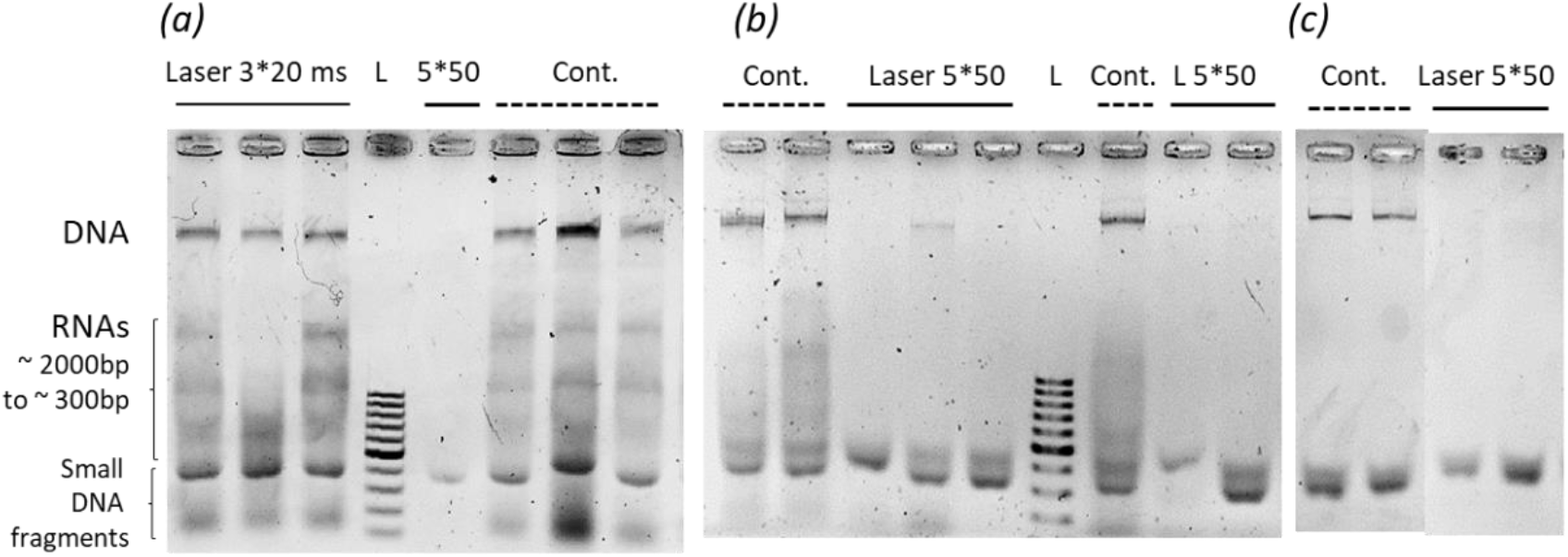
DNA and RNAs integrity in *P. minutus* cystacanths following exposure through the cuticle of infected *G. fossarum*, to a Blue-Diode Laser beam of 1.4W at 450nm for 3 pulses of 20 ms or 5 pulses of 50 ms. Cystacanths were deep frozen in liquid-N just after dissection (less than three hours after laser exposure) (a, c) or after evagination test (1.5 hs at 42°C) (b); (a, b) no RNAse treatment, (c) RNAse treatment after DNA purification and before lyophilisation. L: 500 bp ladder.

The Odds of DNA degradation following laser irradiation at 5*50 ms was 3.5, which is lower than the Odds of evagination failure (24.1). Therefore, some cystacanths may not evaginate but still harbor intact DNA following the 5*50 ms treatment. This discrepancy was even more pronounced in the 3*20 ms treatment where all cystacanths had intact DNA despite an evagination failure rate of 72.7% (N=9 and N=22, respectively).

Laser treatment and its interaction with infection status impacted gammarids’ survival, while infection status alone had no significant effect (Cox multiple regression: N = 632; LR-test = 46.7, df = 5, *P* < 0.0001: laser treatment: Chi^2^ = 28.9, df = 3, *P* < 0.0001; infection status: Chi^2^ = 0.04, df = 1, *P* = 0.84; interaction between infection status and laser treatment, Chi^2^ = 16.7, df = 3, *P* = 0.0008).

Gammarids mortality was higher following irradiation at 5 pulses of 50 ms at 1.4W compared to unexposed controls (z = 4.76, *P* < 0.0001), to gammarids exposed to predrill shot only (1 pulse of 100 ms, 0.5W), and to those exposed to 3 pulses of 20 ms at 1.4W (Fig. 4). The two latter did not differ from unexposed controls (z = 0.10, *P* = 0.91; z = −0.42, *P* = 0.67; respectively). The survival of infected gammarids following laser exposure at 5 pulses of 50 ms at 1.4W was higher than that of uninfected ones (z = −3.53, *P* = 0 0.0005; Fig. 4; plain lines versus dashed lines).

**Figure 4.**
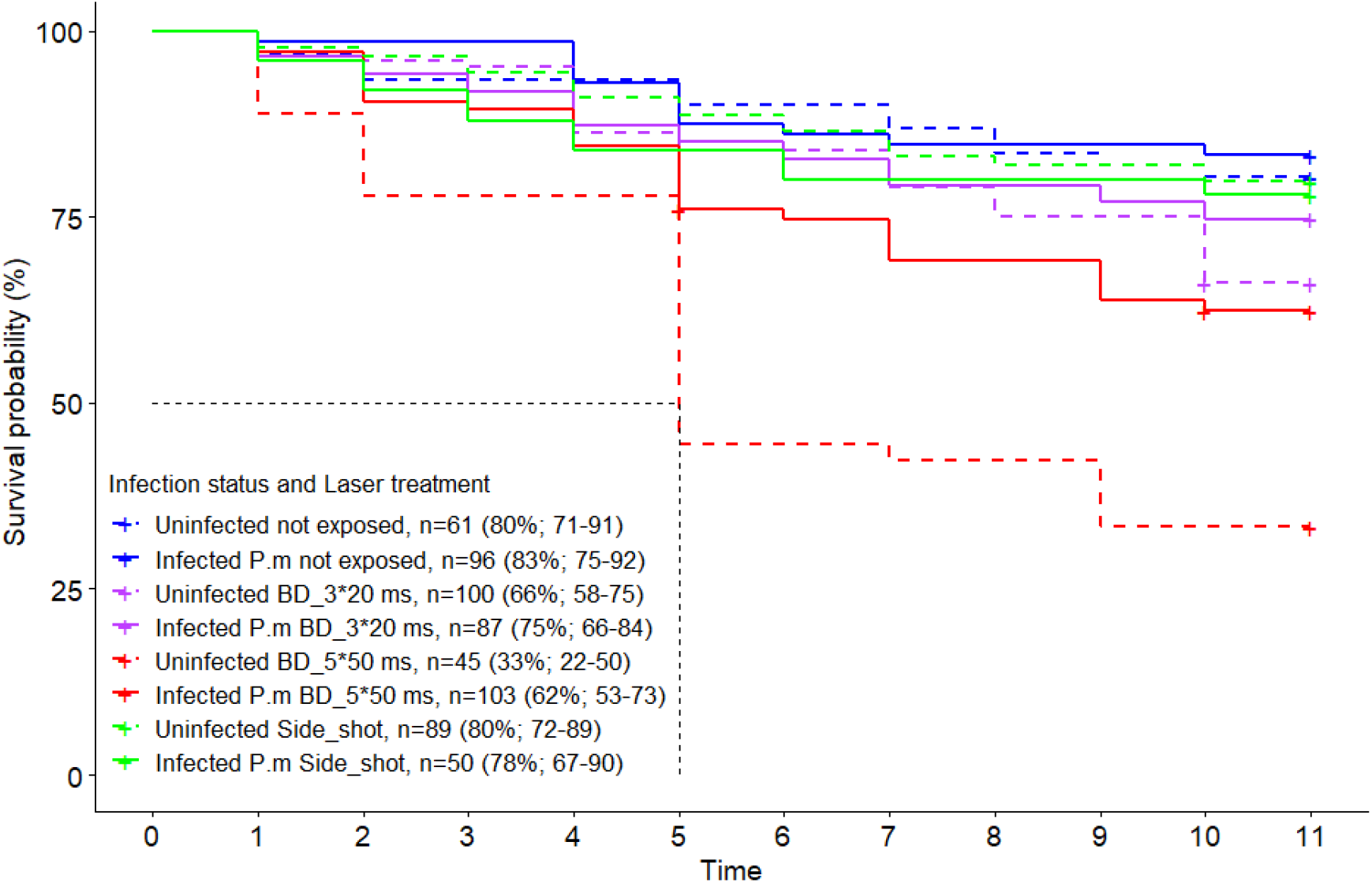
Diagnostic Plot for Cox Proportional Hazards Model representing the survival rate of uninfected gammarids (dashed lines) and gammarids infected with *P. minutus* (plain lines) up to 11 days post-irradiation to Blue Diode laser beam at 450 nm. Gammarids were exposed to one of two treatments: 3 pulses of 20 ms at 1.4 W interspaced by 100 ms (purple lines), or 5 pulses of 50 ms at 1.4 W interspaced by 100 ms (red lines). In both treatments, gammarid’s cuticle was pre-drilled by irradiation at 0.5W, 100 ms. Two controls were processed in the same way (anesthesia and handling) but either not exposed (blue lines) or exposed to a single pulse at 0.5W, 100 ms (next to *P. minutus* for infected ones) (Side_shot). n=Sample size per treatment. (%): Probability of survival at 11 days and associated 95% CI.

The wound left by laser irradiation (enlarged pre-drilled hole) was plugged within 2-3 hs and melanized within 24 hs (pers. obs.), and the scar was removed together with exuviae at the time of molting (Fig. 2).

### Efficiency of laser treatment with the Blue-Diode laser and the Nanoseconds-Green laser on *P. tereticollis*

We tested several protocols with the BDL varying in the number and duration of pulses, to kill *P. tereticollis* while preserving gammarids survival. All *P. tereticollis* cystacanths survived to exposure at 8 * 50 ms, according to DNA integrity assay (N=13; Fig. 5c). Sublethal laser treatments often led to *P. tereticollis* evagination within live gammarids (Fig. 5a, b). All treatments involving 5 to 8 pulses of 100 ms at 1.4W resulted in a rapid decrease in gammarids’ survival (50% within 48 hs, 0 to 10% 11 days after exposure). Most cystacanths exhibited visible wound following exposure to 5 pulses of 100 ms 1.4W (Fig. 5c), and their evagination success rate was decreased to 16.3 % (N=43). Yet, DNA integrity was preserved in 7 out of 10 samples (Fig. 5c).

**Figure 5:**
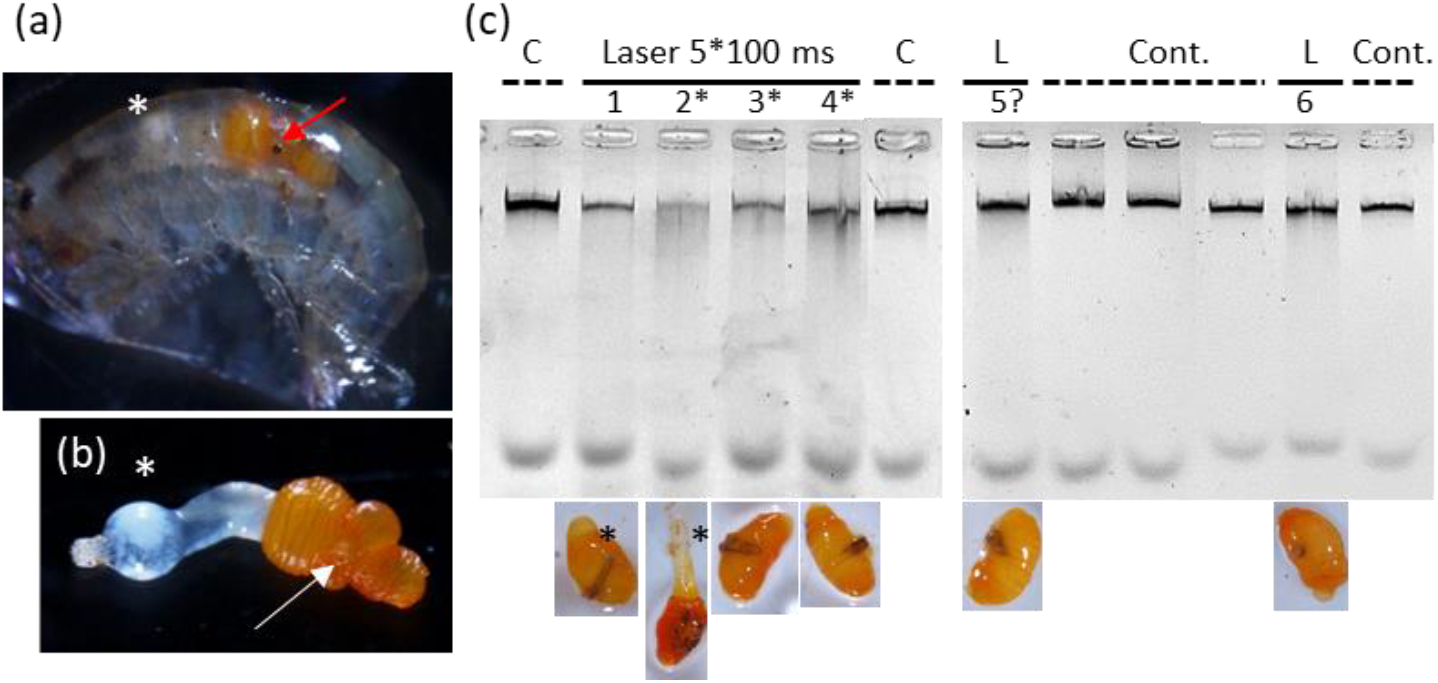
Effect of exposure with Blue Diode Laser on *P. tereticollis:* (a, b) Appearance of the live gammarid host and of *P. tereticollis* cystacanth dissected out 24 hrs after exposure to 3 pulses of 100 ms at 0.5 W, 450 nm: the amphipod is harboring a small wound at the impact point (red arrow) and *P. tereticollis* has evaginated its proboscis within its live host (white tubular structure with a bulb: white asterisk). (c) DNA integrity of *P. tereticollis* cystacanths dissected out from live gammarids two days after exposure to 5 pulses of 100 ms at 1.4 W, 450 nm: C, controls not exposed to laser; L, cystacanths exposed to laser treatment with the corresponding picture below DNA sample. (*) possible partial degradation of DNA (mild smearing relative to the band intensity of high molecular weight DNA). Although cystacanths exhibited a clear wound, and full (lane 2*) or partial (lane 1) evagination of the proboscis, DNA was either partly degraded or intact. DNA purification included RNAse treatment.

The Nanoseconds-Green laser was tested on *P. tereticollis* as an alternative to BDL. We tested the effect of high peak intensity on *P. tereticollis* survival by exposing infected gammarids to 2 shots of 5 pulses, each between 4 and 8 mJ per pulse, 40 Hz. Four to five hours after exposure, all gammarids were dead (N = 10), while 70% of *P. tereticollis* were evaginated within host and still alive, as evidence by their contractions in reaction to light after dissection (N=13). At 4 shots of 5 pulses, all gammarids died and 58% *P. tereticollis* were still alive despite visible wound (N=12) (electronic supplementary material, Fig. S9). We then tested an alternative protocol to increase the deleterious effects of the NGL beam on *P. tereticollis* while decreasing lethal damages to host, based on a “jackhammer”-like effect. A pilot test was run by irradiating *P. tereticollis* infected gammarids at 50 to 100 pulses at 40 Hz, and 2 mJ energy per pulse. After three weeks, all *P. tereticollis* were still alive, as established by evagination test or by observation of evaginated cystacanth within gammarid’s body in reaction to host’s death (N=8).

## Discussion

Laser-based technologies are increasingly used in biology. Here we took advantage of the destructive effects of high-irradiation energy of laser on living organisms to investigate its potential use in de-parasitizing live amphipods from their internal helminth macroparasites. We choose two sources of powerful monochromatic light in the visible spectrum - 450 nm and 532 nm - corresponding to the absorption peak of the parasite’s integumental chromophores (carotenoids) and lipids, respectively. We tested various laser treatments using a newly designed, versatile, and partly automatized full setup, and assessed their efficiency in killing the cystacanths of *P. minutus* and *P. tereticollis* using two criteria to assess parasite’s death: the capacity to evaginate its proboscis and DNA integrity. We successfully optimized a treatment to kill *P. minutus* inside live gammarids using BDL at 1.4W, with 5 pulses of 50 ms, despite non-negligeable mortality of gammarids with a survival probability decreased by 20%.

However, we failed to adapt the technique to the larger acanthocephalan *P. tereticollis:* although exposure to 5 pulses of 100 ms, 1.4W resulted in cystacanth’s evagination rate below 20% and DNA integrity below 60%, gammarids mortality was very high as well (close to 90% after 11 days). Interestingly, *P. minutus* has higher concentration of carotenoids compared to *P. tereticollis* (Perrot-Minnot et al. 2011), therefore the higher efficiency of BDL on this species is consistent with the absorption of laser energy by these chromophores and the consecutive lethal thermal effects. The absorption by *P. minutus* of most of the energy delivered by BDL is further supported by the higher survival of irradiated infected gammarids compared to irradiated uninfected ones, showing the deleterious effects of laser energy when not absorbed by its target. We suspect that both the lower concentration in carotenoids and the larger size of *P. tereticollis* accounted for the failure to induce lethal damages even after increasing energy delivered by the BDL. Indeed, the thermal effects of BDL involved slow continuous heating which could more easily induce lethal damage in the smaller and carotenoid-rich *P. minutus* compared to the larger *P. tereticollis*.

As an alternative to thermal effect only, we used the NGL to induce photo/thermomechanical damages on this acanthocephalan species. However, despite visible wounds, the damages induced to *P. tereticollis* at irradiation levels still compatible with intermediate level of host survival, did not induce instant death. In fact, many cystacanths were found partly or completely evaginated within the host few hours or days after irradiation, adding another source of mortality to gammarids. It is possible that the high-energy and rapid shock wave produced by NGL resulted in limited superficial thermal ablation of parasite’s thick and multilayered body wall, and that average heating was insufficient due to low total energy delivered.

With respect to the assessment of death, we have shown using DNA integrity assay that evagination failure was not reliable evidence for death in these parasites. Evagination success has been used in previous study addressing the protective function of carotenoid accumulation against UV and the consequence of UV exposure on parasite’s ability to establish in its final host as a proxy for parasite’s fitness (Perrot-Minnot et al. 2011). Here however, we needed a diagnostic assay for parasite death, i.e., the definitive and total disruption of parasite-host interactions. The integrity of DNA thus appeared as a reliable assay to assess parasite death (Roy et al. 2020).

Gammarids mortality was observed with both laser sources but was much higher with the NGL. This is consistent with the two distinct effects of these laser sources. The BDL possibly produced a mild thermal effect that did not abruptly damage vital organs but rather induced limited heating over a short time (possibly attenuated by exposing gammarids to laser in a low volume of refrigerated water). By contrast, the NGL produced a short-in-time mechanical wave at high energy, which could more likely irreversibly damage surrounding host’s cells. There could also be multiple causes of gammarids death apart from direct/immediate effects. During the first four days following exposure to the optimal BDL treatment to kill *P. minutus*, mortality was low and comparable to that of gammarids exposed to the “pre-drilled” treatment (one pulse of 0.5W, 100 ms) while slightly lower than controls. In *P. minutus*-infected gammarids, the clotting and melanization of laser wound occurred within few hours and the scar left on the cuticle at the laser impact was fully melanized within 24 hs, a time window comparable to wounds produced mechanically with a thin needle (MJPM, pers. observation). If hemolymph loss was responsible for impaired survival to laser treatment, mortality of pre-drilled gammarids and of gammarids exposed to the full treatment would have been comparable, providing that hemolymph loss was comparable between these two groups. Alternatively, three non-exclusive indirect effects of laser treatment may account for this delayed mortality of infected gammarids: persistent cellular damages due to thermal diffusion of residual energy, the physiological cost of recovery (for instance to replenish hemocytes), or the excretion of products by the dying parasite. The high mortality of gammarids following laser irradiation of *P. tereticollis* could be due to two non-exclusive consequences of high-energy laser light: collateral damages to host cells due to diffusion, and the parasite reaction to stress inside its host, mainly evagination. Cystacanth’s evagination could have been either the cause or the consequence of gammarids death and both were observed (Fig. 6). High mortality with NGL could result from irreversible damages to cells, in particular blood cells responsible for clotting and wound melanization. Indeed, none of the dead gammarids shown sign of clotting or melanization at the impact point few hours after irradiation with NGL, in contrast to those exposed to the BDL. Photomechanical treatment may have specifically damaged cells by the shock wave, causing gammarid’s death even if the total energy delivered is low.

Killing acanthocephalan cystacanth inside its live intermediate host will allow us to evaluate the reversibility of phenotypic manipulation by *P. minutus*. The same goal applies to *P. tereticollis* providing that laser treatment can be optimized to minimize the death of *G. fossarum* associated with higher energy irradiation. To go further, the development and application of laser-based de-parasitization could also be expended to other host-macroparasite systems with various physical characteristics, such as parasitoids inside their insect hosts or insect larvae inside plant seeds. Indeed, the various thermomechanical and thermal effects of high-energy, short-wavelength visible light can be exploited and adapted to a range of host-parasite systems, providing that an appropriate imaging and aiming system allows targeting the microparasite inside its living host. Such sighting system includes direct visualization, as used in the present study, but also X-ray scanner or micro-computed tomography (O’Sullivan et al. 2018). The temporal and spectral characteristics of the lasers as well as the irradiance and the tempo of the beam should then be optimized to choose the laser-tissue interaction modalities best suited to the types of pigments or physical structures that must absorb the energy delivered, while minimizing damage to the host’s surrounding tissues.

## Supporting information

Supplementary material from "Laser-based killing of a macroparasite inside its live hosts"

## Acknowledgements

We would like to thank Baptiste Paulmier for his help with software development, and Jean-Emmanuel Rollin for help with gammarids sampling. We thank Dr Frédéric Marin for giving access to the freeze-dryer used for DNA concentration.

## Data accessibility and supplementary information availability

Raw data are available in Mendeley Data. DOI: 10.17632/8j78n2znjb.1 Supplementary material are available online.

## Authors’ contributions

OM and MJPM together conceived the idea for the project, formulated the research goals and adapted the design during the study. OM modelized, designed and assembled the full setup, and designed the software. OM, MJPM, and AB performed the experiments. MJPM processed and analyzed the data. OM and MJPM equally contributed to drafting, editing, and finalizing the manuscript.

