## Supplementary material from "Laser-based killing of a macroparasite inside its live hosts" for "Laser-based killing of a macroparasite inside its live invertebrate host"

S1 Background general information on laser-material interactions, laser beam specification and optical quantities

S2 Laser sources specification and the design of full setup

S3 Parasite death: DNA integrity assay

### S1 Background general information on laser-material interactions, laser beam specification and optical quantities

#### *General information on laser - material interactions usable for de-parasitization*

Depending on the type of laser used, it is possible to obtain very variable effects on the irradiated material. These effects can be thermal, such as heating, fusion or vaporization, photochemical - molecular denaturation - or photomechanical such as ablation, leading to a more or less violent expulsion of material from the exposed substrate. In our study, the effect used was essentially thermal, but differs between the two laser sources tested by their kinematics and induced effects. These effects correspond to the appearance of a thermal gradient of variable duration, and possibly one or more phase changes in the material. All are based on optical absorption, and on the local transformation of absorbed light into heat following Beer-Lambert's law. In general, the laws of geometric optics are no longer applicable laser beam behavior, but rather the model of Gaussian beams. This model describes the distribution of light intensity in the spatial profile of a laser beam, and laser beam behavior during its propagation and more practically when it is focused by a lens. Contrary to popular belief, all laser beams are divergent. Then they are often defined by the value of their divergence and their waist.

#### *Laser sources and their optical shaping systems: generalities*

The usual and simple distribution of energy or power in a laser spot follows the equation (1).

$$I(r,t) = I_0(t).e^{-2.(\frac{r^2}{\omega_0^2})} \qquad (1)$$

This equation being worth zero only for r tending to infinity, the radius of a laser beam is therefore defined by the value $\omega_0$, called beam waist (2).

$$I(\omega_0,t) = I_0(t)/e^2 \qquad (2)$$

It also means that the energy or power contained in a circle of radius r = $\omega_0$ represents 86.5% of the total energy or power. Figure S1 presents an example of gaussian laser beam.

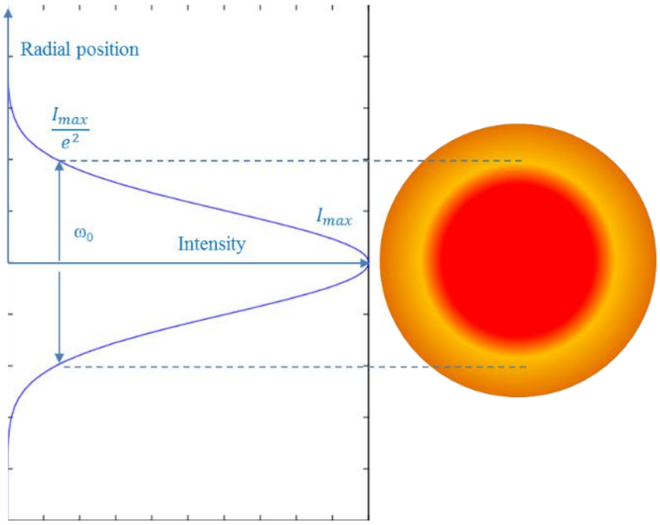


Figure S1 : Intensity distribution in a gaussian laser beam and definition of the beam waist $\omega_0$

The equation (1) describes a fundamental transverse mode laser beam ($TEM_{00}$) comparable to the best possible laser beam for a given laser source. In general, the laser beam used is degraded compared to this ideal solution, it is larger and more divergent than the $TEM_{00}$ mode. To account for this degradation, a beam-quality factor, $M^2$, is quantified. The $M^2$ factor of a laser beam of wavelength λ is estimated as the ratio between the beam product of the available laser beam - defined as the product of waist multiplied by divergence - and the best laser beam that could be obtained (the mode $TEM_{00}$). If $M^2 = 1$, the laser beam is said to be single mode transverse, and if it is greater than 1 it is said to be multimode transverse. The $M^2$ factor have a large impact on the propagation of a laser beam and more specifically on its ability to focus on a small area with a large Depth Of Focus (DOF).

If $\omega_0$ is the smallest radius of a laser beam of quality $M^2$ in the focal plane of a converging lens with focal distance f, the expression of the beam radius as a function of the distance from this focal plane $z_0$ will be expressed with the equation (3)

$$\omega^2(z) = \omega_0^2 + \left(M^2\right)^2 \left(\frac{\lambda}{\pi\omega_0}\right)^2 (z - z_0)^2 \text{ with } \omega^2(z_0) = \omega_0^2 \qquad (3)$$

The propagation of the envelope in $I_0/e^2$ will therefore follow that presented in Figure S2.

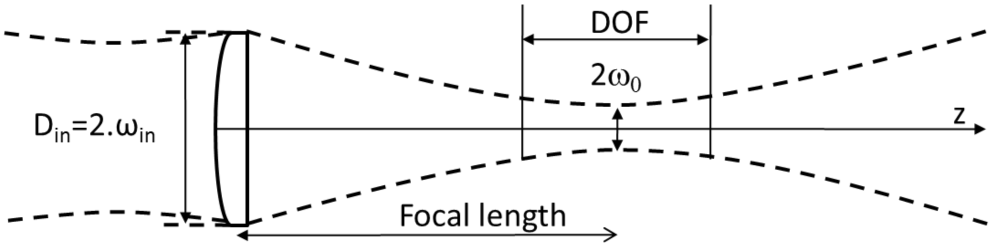


Figure S2: Illustration of laser beam propagation after passing through a focusing lens

In general, a focusing lens is used to increase laser intensity on the sample, and the sample is positioned in the focal plane of the lens. If the thickness of the sample subjected to the incident laser beam is small, laser beam diameter can be considered initially as constant. The diameter $d_0$ of the laser spot in the focal plane is then calculated using equation (4) where $D_{in}$ is the diameter of the laser beam at the lens entrance:

$$d_0 = 2.\omega_0 = M^2 \frac{4\lambda f}{\pi D_{in}} \qquad (4)$$

Laser spot diameter varies around this smallest observable value in the focal plane of converging lens. As in geometric optics, it is also common to define a collimation length called Rayleigh distance in Gaussian optics, and denoted $Z_R$. The parameter $Z_R$ is such that laser spot diameter changes by a factor $\sqrt{2}$ over this distance, and therefore laser spot area is multiplied by 2 and irradiance is halved. This variation is therefore very important, and we prefer to use an additional parameter to determine the depth of focus (DOF) on each side of the focal plane for which laser beam diameter increases from 1 to 1+ k (equation 5).

$$DOF = 2 \times z_k = 2 \times \left(\frac{\lambda M^2 f^2}{\pi\omega_{in}^2}\right)\sqrt{\left(k^2 + 2k\right)} \qquad (5)$$

With our experimental setup, the maximum diameter of the laser spot of a beam of $M^2 = 2$ @ 532 nm (maximum value coming from equation (7) and table (1)) focused by a converging lens of focal length 100mm and with a 10 mm entry diameter in front of the lens, gives a minimum spot diameter of:

$$d_0 = M^2 \frac{4\lambda f}{\pi D_{in}} = 2 \times \frac{4 \times 532.10^{-9} \times 0.1}{\pi \times 5.10^{-3}} \cong 30\mu m$$

The DOF distance for a maximum variation of + 20% on the diameter of the laser spot (i.e. + 40% on the area of the laser spot and therefore -40% on the irradiance) will be:

$$DOF = 2 \times \left(\frac{\lambda M^2}{\pi}\right)\left(\frac{f}{\omega_{in}}\right)^2 \sqrt{(k^2 + 2k)} = 2 \times \frac{532.10^{-9} \times 2}{\pi}\left(\frac{0,1}{2.5.10^{-3}}\right)^2 \sqrt{0.2^2 + 0.4}$$
$$\cong 0.8mm$$

It will therefore be necessary to adapt the optical assembly according to the laser sources used, their wavelengths and their beam qualities.

This deposited optical energy is almost instantly converted into heat. Induced heating is then distributed temporally and spatially in the material according to its thermal properties, for example conductivity or thermal diffusivity. In general, thermal effects can be divided into two main categories according to laser exposure time: (1) slow effects, where heat generation time and propagation time are of the same order of magnitude; (2) rapid effects, which correspond to an energy supply in the material and its transformation into heat almost instantly compared with the duration of heat propagation. The two tested laser sources correspond to these two categories, and can be summarized as follows: (1) The use of a continuous laser source with output power around the watt, possibly chopped, with an exposure time of the order of a second to a few seconds; (2) The use of a pulse laser source with a very short pulse duration typically in the nanosecond range (<100ns) with low pulse energy (<10mJ) and a low repetition frequency (<100Hz).

Whatever the laser source used, the amount of light entering the material is that which has not been reflected or backscattered at the surface or at the interface of media of various types. This amount of remaining energy is likely to be transformed into heat via absorption. Depending on the amount and time of heat production, this transformation will lead to variable temperature rise and gradient. The irradiated area will then be susceptible to transformation. This area called HAZ (Heat Affected Zone) will not be homogeneous because the material will be specifically transformed according to the temperature reached locally and the duration of exposure to this temperature. It will then be possible to heat, melt or vaporize a material but also to coagulate, carbonize or even fragment it. A detailed description and examples can be found in [1-4].

***Thermal Laser - tissues interaction***

Exclusively thermal effects require longer exposure time, for example to laser source with continuous emission or long pulse duration. The temperature reached corresponds to the balance between the amount of heat provided by laser light absorption and the amount of heat escaping by thermal conduction. In addition to heat loss, energy is also used to either achieve a phase change or chemically transform the tissues, for example by carbonization [1-

4]. However, the temperatures reached locally are much lower and the processes much slower, we can therefore neglect induced mechanical effects.

Thermomechanical effects are initiated by the absorption of part of the incident light from a pulse of short duration, typically from one to a few tens of nanoseconds. During absorption, peak energy illumination at the impact zone is very high and the relaxation of the energy absorbed in the form of heat is extremely rapid. This relaxation then leads to a very rapid and much localized heating of the material. As the temperature increases very quickly, the naturally rather slow thermal conduction and diffusion phenomena are almost negligible. This then causes a localized and almost instantaneous phase change of the material, initially solid or liquid, in the form of a vapor bubble. This vapor continues to absorb the laser radiation and can go as far as being at least partially ionized; this ionized vapor is then called plasma. The vapor or plasma thus formed relaxes and leads to the generation of a shock wave capable of mechanically damaging the irradiated tissues, for example by inducing ablation or fractures. In general, the amount of ablated material depends on laser parameters, lighting conditions, the focusing on the sample, and the sample's chemical composition and structure. Each laser shot induces an ablation of a certain depth. By multiplying laser shots, it is then possible to dig the sample. Note that it is not compulsory to work in nanosecond mode to generate laser ablation: it is also possible to obtain a purely "thermal" cavitation (for example by using a microsecond source at 2 or 3 µm that is at the peak of water absorption, and violently relaxing the bubble of steam generated). This technique is used for example in laser lithotripsy. It is also possible to create this ablation by non-thermal photo-ablative process (using a nanosecond UV laser for example). However, if the laser impacts are temporally close, slow thermal effects are no longer negligible and part of the heat produced by each laser pulse tends to accumulate over laser shots. It will then be necessary to consider the slow thermal effects as described above, in addition to the photomechanical effects.

#### *Definition of optical quantities*

The optical energy quantities characterizing the light coming from laser sources are specified below:

- The Radiant Energy Ep usually named light energy; it can be described as the energy contained in a pulse of power P and duration Δt; the SI unit (International System of Units) is the Joule (J).

$$E_p = \int_0^{\Delta t} P(t) \cdot dt \qquad (6)$$

- The peak radiant flux or peak power; the SI unit is the Watt (W)

$$P_{peak} = \frac{E_p}{\Delta t} \qquad (7)$$

- The average radiant flux or average power, in Watt (W), with F the repetition rate in Hertz (Hz)

$$P_{average} = E_p . F \qquad (8)$$

- The irradiance or flux density, usually named intensity; SI unit is the Watt / area (W.cm$^{-2}$).

$$I_{peak} = \frac{P_{peak}}{(\pi \cdot \omega)^2} \qquad I_{average} = \frac{P_{average}}{(\pi \cdot \omega)^2} \qquad (9)$$

### S2 Laser sources, specification, and design of the full setup

#### *Blue diode laser source*

Several assemblies based on laser diodes emitting in the blue at around 450nm have been built and tested. These uses either a low power laser diode 100mW @ 450nm (PL-B450 OSRAM®) with a transverse single mode beam ($M^2$ = 1) or a multimode laser diode capable of emitting 1.6W @ 450nm (PL-TB450B OSRAM®) but in multimode with an $M^2$ of about 4. Only the latter is described here because it is the one that has been chosen based on preliminary tests. As desired, this laser diode has its emission wavelength centered on the absorption peak of β-carotene [4]. The laser diode is very divergent and astigmatic, with a large difference of divergence in the two directions orthogonal to the direction of propagation. To be able to generate a small laser spot of diameter <100µm on the specimen, it is therefore necessary to transform the initial beam by adding optics. The laser beam shaping uses a first aspheric collimating lens with short focal length (Thorlabs / Geltech-Lightpath C971TME-A) associated with a unidirectional telescope allowing for obtaining a quasi-circular beam (Thorlabs LK1087L2.1 and LJ1402L1-A1). This beam is then focused on the sample via a 100mm focal achromat (AC254-100-A-ML1). The assembly is simulated on Zemax OpticStudio® software to optimize the inter-element distances and then designed using Autodesk Inventor®. It is then machined in Fortal alloy or copper for the laser diode mount, and then all optical components are glued. A diaphragm is placed along the rapid axis to reduce geometric aberrations.

Energy distribution at the focal point is presented in Fig. S3. At the output of the assembly in the focal plane, we obtain a power of 1.4W from the 1.6W of the laser diode. The minimum spot size is f 70 * 30 µm corresponding to an irradiance of 67kW/cm$^2$.

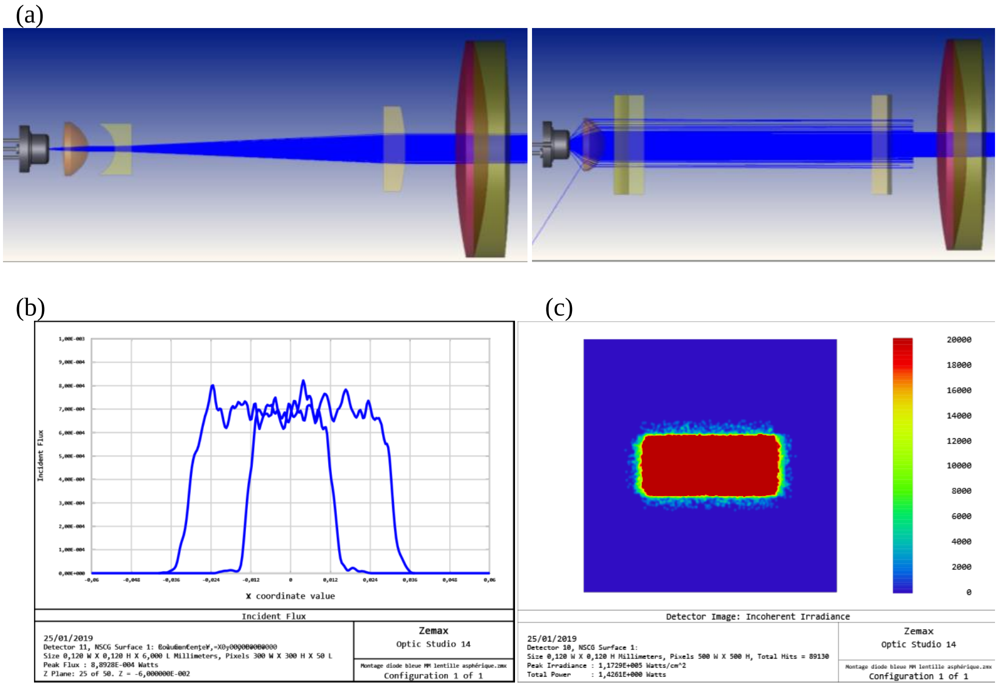


Figure S3: (a) Zemax views of the complete optical assembly in the two main directions, slow axe, and fast axe; (b, c) X and Y beam profiles in the focal plane for the two directions (fast and slow axes)

Variation in laser spot diameter is calculated around the focal plane for the two main axes X and Y and over approximately ± 2mm i.e., over a depth of field (DOF) of approximately 4mm maximum (Fig. S4):

- On a DOF of ± 1mm, laser spot dimension ranges from 64 to 81µm and from 28 to 45µm
- On a DOF of ± 2mm, laser spot dimension ranges from 64 to 105µm and from 28 to 65µm

The corresponding energy illumination ranges from 77kW/cm$^2$ to 39kW/cm$^2$ (~ -50%) and from 77kW/cm$^2$ to 20kW/cm$^2$ (~ -75%), for DOF of $\cong \pm 1mm$ and$\cong \pm 2mm$ , respectively.

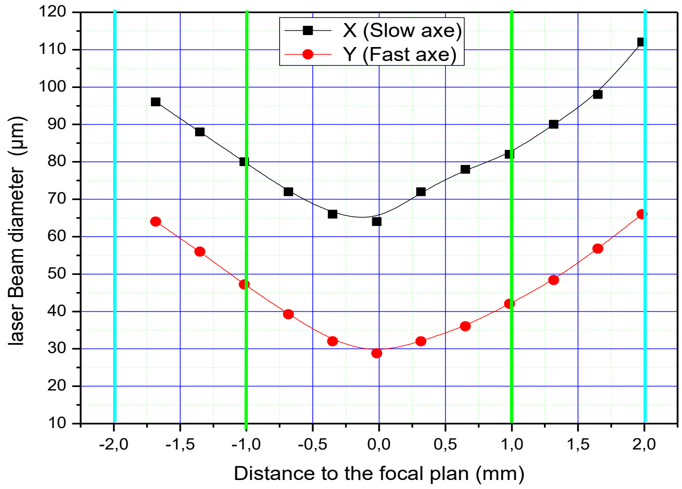


Figure S4: Laser spot diameter in 1 / e$^2$ around the focal point for the two main directions X and Y, according to the distance to the focal plan.

The Blue Laser Diodes (BLD) are controlled by a power supply developed in the laboratory. This driver includes a color touch screen coupled to an Arduino Due which drive two OEM blue laser diode power supplies from Thorlabs® (IP250-8V and LD3000R). These two power supplies allow current and power control of laser diodes from 0 to 2.5A (multimode transverse diode 1.6W@450nm), with the sufficient output voltage needed for blue laser diodes. The controller is presented with its visual interface in Figure S5. Several menus allow the user to configure the driver and choose the desired operating mode as continuous or pulsed. It is for example possible to program a low aiming current (visible by the camera but without damage to the specimen) and to configure sequences of light shots by adjusting the ON times, the OFF times, the current of each pulse and their number.

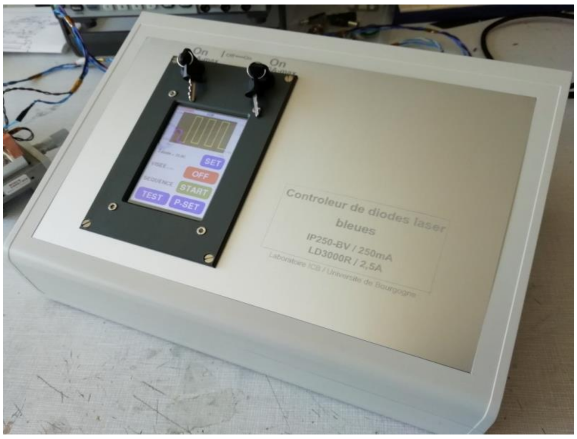


Figure S5: Picture of the Blue Diode Laser driver, and example of a multi pulse configuration

***Nanosecond green laser source***

The nanosecond laser is a laser source developed in the laboratory initially for laser-induced plasma spectroscopy (LIBS) application. This laser is based on an Nd: YAG crystal pumped longitudinally by a quasi-continuous laser diode stack. It is actively triggered by an electro-optical Q-switch cell which makes it possible to obtain pulses of short durations, around ten nanoseconds. Its natural emission wavelength at 1064nm, i.e., in the near infrared is then converted into green emission at 532nm, thanks to a frequency doubling stage based on a KTP crystal. The performances of this source are:
- Adjustable output energy from 1 to 12 mJ @ 532nm
- Pulse duration around 10ns (depends on pumping and therefore on the energy extracted)
- Repetition frequency from 1 to 40Hz
- $M^2$ beam quality <1.5 over the operating range
- Shot-to-shot stability at 2σ <2%

A computer driver including a touch screen makes it possible to adjust all the laser's parameters, such as firing frequency, the number of pulses and energy per pulse. The performances obtained at laser output are measured by dedicated metrology devices: for energy measurement, an OPHIR® Nova controller and a PE25BB pyroelectric head, for the pulse duration, a fast photodiode DET10A – Thorlabs® associated with a fast oscilloscope Tektronix®TDS5052B / 500MHz-5Gs / s.

A vertical plate containing the different optical functions is placed at the exit of the laser, allowing the focusing of the beam on the specimens placed on the specimen holder. This plate includes:
- An X2 magnification telescope allowing laser spot diameter in the focusing plane to be reduced by twice.
- A laser-aiming diode emitting in the red near 630nm, with a diaphragm placed at the output in order to limit the laser spot to a well circular point of small dimension, far below the average size of a parasite.
- Adjusting mirrors making it possible to superimpose the red aiming beam and the green power laser beam.
- A 100mm focal focusing lens mounted on a manual displacement micro stage, to properly position the focal plane at the level of the specimen.

The plate and all the mechanical elements of the assembly are designed using Autodesk Inventor and manufactured in Fortal alloy by 3-axis and 5-axis machines in the mechanical workshop (CRM) of ICB laboratory.

The 100mm converging lens makes it possible to obtain a laser spot ≤50µm in diameter corresponding to a fluence of a few tens to a few hundred $J.cm^{-2}$, and a peak illumination of up to several $GW.cm^{-2}$. In this field of illumination, the interactions are very photomechanical with the possible appearance of a plasma.

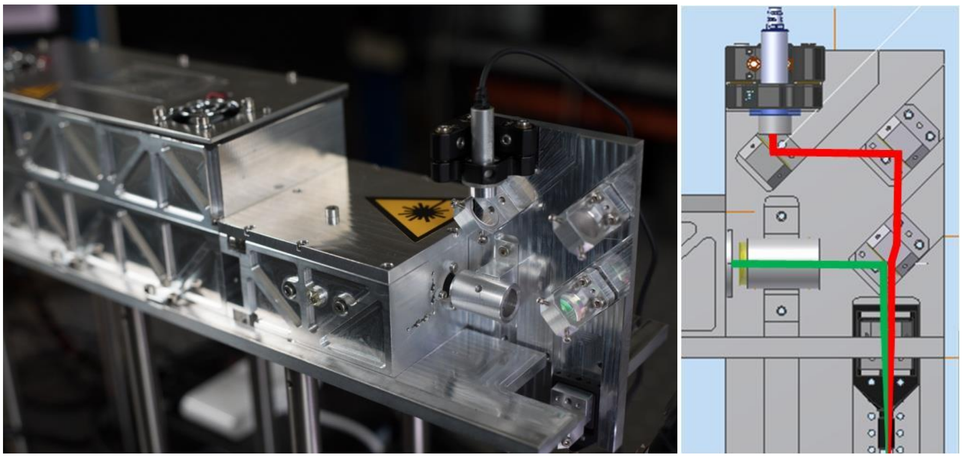

Figure S6: Nanosecond laser source with focusing head

***Thermoregulated sample holder***

Gammarids are freshwater crustaceans of small size (approximately 4mm to 8 mm body length). They can reach high level of locomotor activity when they are stressed, for example under strong light or with a touch stimulus (escape response). To irradiate the parasite inside its live host while limiting the damages to surrounding host tissues, it is necessary to precisely position the laser impact point on the parasite. This can be achieved by complete immobilization of the gammarid under anesthesia, in a suitable refrigerated sample holder.

We designed a sample holder with the following features:

- Cool the specimen to a stable, reproducible, and adjustable temperature from 0 to 20 ° C, with a target temperature of 3 to 4 ° C.
- Facilitate the handling of gammarus (manual placement and removal)
- Can be cleaned and disinfected with alcohol.
- Be open to the outside to allow lighting, video viewing, aiming and laser exposure.
- Have a device that is compact and light enough to be adapted to an XY motorized micrometric movement.

The sample holder is made up of several elements mounted one on top of the other and integrated into a mounting block, which is itself fixed to the displacement system. Figure S7 shows the separate elements and the integrated block mounted on the XY movement.
The upper part is machined from stainless steel and incorporates the housings (circular or linear) where the specimen will be placed. This part is then mounted on a large Peltier thermoelectric module (50*50mm). A thermoelectric module is an electronic component that allows heat to be transferred from one "cold" side to the other "hot" side by passing an electric current. Depending on the intensity and the direction of flow of this current, it is therefore possible to regulate the temperature of an object placed for example on the cold face. This module is then mounted on a small copper water heat exchanger. This exchanger removes the heat extracted from the hot face and the heat produced by the thermoelectric module itself. Indeed, the efficiency of these modules is generally at most 50% and it decreases quickly when the temperature difference between the two faces is large (here> 20 ° C). In addition, the cold face must be accessible and visible to place the specimen, illuminate and visualize them with the camera and of course expose them to the laser. It is

not possible to thermally isolate it, so it is in direct contact with the ambient air. These constraints of low efficiency and the need for a compact and light assembly therefore lead to the choice of a water exchanger instead of an air exchanger, which would have been too bulky and ineffective. The assembly is then placed in a part machined in Fortal® and blocked by a nylon cover which makes it possible to maintain the whole and to fix the assembly to the displacement system. Two NTC thermistors (Negative Temperature Coefficient resistors), used as temperature probes, are glued to the conductive adhesive on the upper part; one probe is used for regulation and the second for a control measurement.

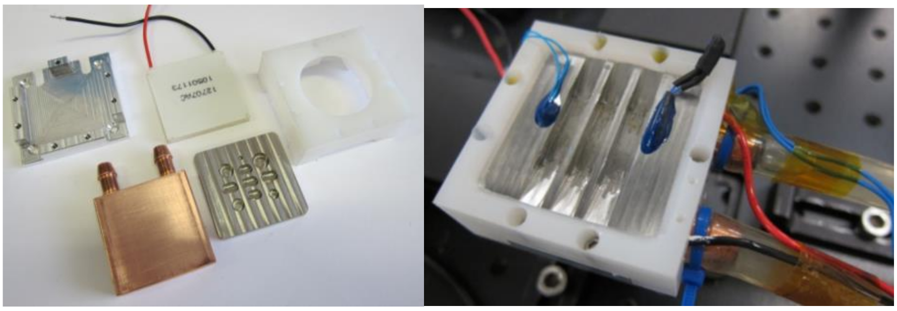

Figure S7: Views of the sample holder, elementary parts on the left and full device mounted on the right. Two types of specimen holder are presented: with circular housing on the right and with linear channels on the right

The sample holder is connected to a box comprising the electrical power supplies, a water / air heat exchanger, and a water pump. Control and regulation are based on an Arduino Due module and a color touch screen, which are integrated in a remote console. The configuration and control of temperature, including graphic output, is depicted in Figure S8. Programmable temperatures range from -10 ° to + 40 °. For regulation at 3.5 ° C for example, the time required to reach this temperature is less than a minute and the stability obtained is ± 0.1 °C.

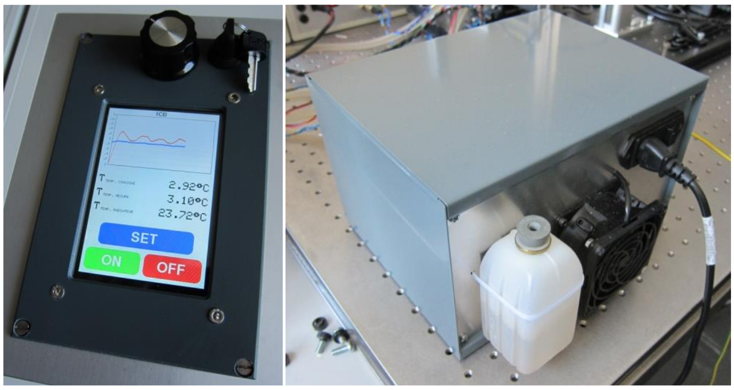


Figure S8: Cooler box and control

#### *Full setup*

The laser sources are mounted on an optical table so that the laser beam is directed vertically from top to bottom. The nanosecond source and its beam shaping system are mounted on a high platform held by six 200mm rods. The blue laser sources are mounted on three 40mm-side Microcontrol® rails for easy z-adjustment to the specimen. A color camera with white LEDs lighting (Dino-Lite® Premier) is mounted on a vertical translation system to optimize the focusing on specimen, the camera's optical axis then makes an angle of about 20 ° from the optical axis of the lasers. The XY motorized translation system and thermalized sample holder are placed in the focal plane of lasers and camera. The micrometric positioning system chosen is based on the use of commercial Thorlabs® components: two motorized displacements (Thorlabs® MTS25-Z8) each associated with a controller (Thorlabs® KDC101). The two displacements are mounted orthogonally to each other to cover an area of 25*25 $mm^2$ with an accuracy of 1 µm in position. The optical axes of the two laser systems are fixed and separated , and the transition from one system to another is done by moving the XY positioning system between two positions marked on the optical table. Only a camera adjustment is necessary, the positioning systems as well as the display are controlled by two PC software supplied by their own manufacturers.

### S3 Parasite death: DNA integrity assay

We quantified DNA integrity by visualization of total DNA on agarose gel. DNA purification from single cystacanth was restricted to digestion without solvent extraction and subsequent precipitation, as these steps could decrease extraction yield and are not required for the analysis of DNA fragmentation.

Individual cystacanths were deep-frozen in liquid nitrogen upon dissection and stored at -80°C. Cystacanths were grinded in 30 µL of pre-heated Queen's lysis buffer + 5 µL of proteinase K solution (20 mg/mL) using a ball mill (RETSCH MM 400 Mixer Mill) for three rounds of 1 min. at 20 Hz (120.min-1), interspersed with few minutes at 37 - 50°C. Two types of balls were mixed to improve grinding: 3mm diameter stainless-steel balls (RETSCH 22.455.0002) and 0.50.-.0.75 mm diameter glass beads (RETSCH 22.222.004) and incubated for 20 min. The samples were then incubated overnight at 55°C. For a subset of samples, RNAse treatment was performed by the addition of 5 µL of RNAse (10 $mg.mL^{-1}$, Thermo-Fisher) and additional incubation for 30 min. at 37°C before overnight digestion. After centrifugation at 5°C 2000 rpm for 5 min., the supernatants were stored at -80°C. To improve DNA visualization, we performed DNA concentration prior to gel electrophoresis. Samples were freeze-dried (Telstar Cryodos 80) and resuspended in 5 µL of ddH2O. DNA was then visualized after gel electrophoresis in 2% agarose (10 min. at 50V and 20 min. at 100V) and staining in Ethidium bromide.

**S4 Preliminary data on an alternative laser treatment: nanosecond-green laser targeting lipids in *Pomphorhynhus tereticollis*.**

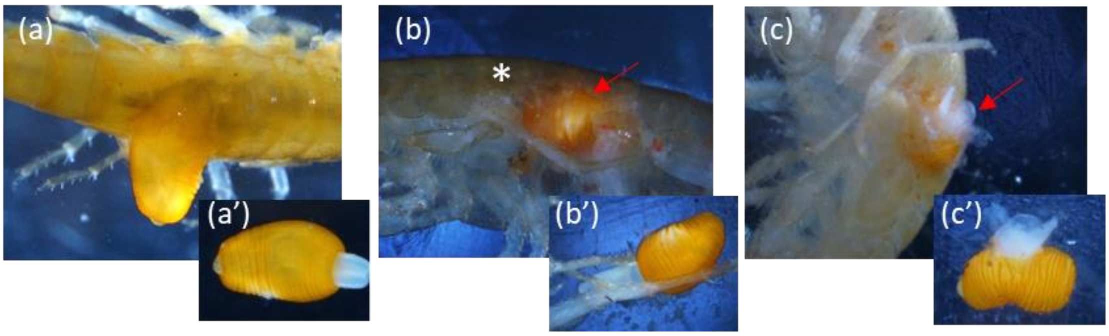


Fig. S9: Appearance of the amphipod host and acanthocephalan cystacanths after exposure to the Nanosecond-Green Laser at 532 nm: partly (a') or fully (b') evaginated *P. tereticollis* cystacanth within (b) or partly outside (a) dead host, following 2 shoots of 5 pulses at 40 Hz; (c, c') damaged (dead?) *P. tereticollis* cystacanth following 4 shoots of 5 pulses at 40 Hz; illustrating tissue ablation/extrusion resulting from the photomechanical effect of nanosecond green laser.
